# MSL2 targets histone genes in *Drosophila virilis*

**DOI:** 10.1101/2022.12.14.520423

**Authors:** Mellisa Xie, Lauren J. Hodkinson, H. Skye Comstra, Pamela P. Diaz-Saldana, Hannah E. Gilbonio, Julia L. Gross, Robert M. Chavez, Gwyn L. Puckett, Tsutomu Aoki, Paul Schedl, Leila E. Rieder

## Abstract

Histone genes are amongst the most evolutionary conserved in eukaryotic genomes, yet *cis*-regulatory mechanisms of histone gene regulation differ considerably amongst species. In *Drosophila melanogaster*, an interaction between GA-rich *cis* elements in the *H3/H4* promoter and the GA-binding transcription factor CLAMP is important for promoting histone gene regulation and factor recruitment to the locus. CLAMP also participates in male dosage compensation by recruiting the Male Specific Lethal Complex (MSLc) to the X-chromosome. We discovered that the male-specific protein of MSLc, MSL2, is recruited to the autosomal major histone locus in *D. virilis* but not to the minor locus or to the single histone locus in other species. While the histone coding sequences are well conserved between species, the critical GA-rich *cis* elements in the *H3/H4* promoter are poorly conserved between *D. melanogaster* and *D. virilis*. We show that CLAMP still targets the two *D. virilis* histone loci *in vivo*. Further, CLAMP interacts with the *D. virilis H3/H4* promoter *in vitro*, even when the poorly-conserved GA-rich *cis* elements are deleted, indicating that the protein interacts differently with the *D. virilis* promoter than it does with the *D. melanogaster* promoter. Since CLAMP and MSL2 directly interact in *D. melanogaster*, we propose that *D. virilis* CLAMP recruits MSL2 to an ectopic autosomal site through interaction with X-like *cis* elements. Further, localization of MSL2 to one of the *D. virilis* histone loci suggests that the loci are regulated differently and that males and females have different requirements for histone gene regulation.

## Introduction

Histones are critical organizational components of eukaryotic chromatin and are highly conserved. For example, histone H3 is 80% identical at the nucleotide level and 99% identical at the protein level between *Drosophila melanogaster* and humans. Histone levels are carefully controlled during both the cell cycle and development; coordinated expression of histone genes is cell-cycle regulated and peaks during S phase (Marzluff *et al*. 2008). Misregulation of histone genes disrupts the precise cell cycle timing during animal development (Amodeo *et al*. 2015; Chari *et al*. 2019). Unsurprisingly, cell cycle regulatory requirements of the replication-dependent histone genes are similar between species (Mariño-Ramírez *et al*. 2006). Despite similar cell cycle and developmental regulatory requirements between animals, histone gene *cis*-regulatory mechanisms appear to differ between species (Kremer and Hennig 1990; Mariño-Ramírez *et al*. 2006).

Organization of the histone genes within the genome also varies widely between species. Vertebrate genomes tend to have lower histone gene copy number and dispersed distribution, while invertebrate genomes carry high numbers of tandem histone repeats. The human genome has two loose clusters of histone genes interspersed with non-histone genes (Marzluff *et al*. 2002). The *C. elegans* genome carries eleven dispersed clusters of the four core histone genes, and histone loci do not include the histone *H1* gene. The histone locus of *Drosophila melanogaster* is a single locus that carries ~100 copies of a histone gene array that includes all five replication-dependent histone genes (Lifton *et al*. 1978; McKay *et al*. 2015; Bongartz and Schloissnig 2019).

The organization and number of histone genes are not well conserved even within Drosophilidae. *D. hydei* has only about 10 histone array copies, and they are located in the middle of euchromatin on Chromosome 4 (Fitch *et al*. 1990). *D. virilis*, which diverged from *D. melanogaster* ~ 40 MYa (Russo *et al*. 1995) has both regular quartet arrays that include the core histone genes (*H2A, H2B, H3*, and *H4*) and polymorphic quintet arrays that include the histone *H1* gene. The quartet arrays are tandemly distributed and linked to a single “major” locus, while the quintets are distributed between both “major” and “minor” loci (Schienman *et al*. 1998).

The diversity in histone gene organization is striking given similar requirements for histone gene expression across species (Mariño-Ramírez *et al*. 2006). In animals, replication-dependent histone biogenesis is controlled by a suite of factors that target histone genes called the Histone Locus Body (HLB) (Liu *et al*. 2006; Nizami *et al*. 2010; Duronio and Marzluff 2017). The interaction between the scaffolding protein Multi-sex combs (Mxc; *Drosophila* homolog of human nuclear protein of the ataxia telangiectasia mutated locus/NPAT) and the RNA processing factor FLICE-associated huge protein (FLASH) is required for HLB formation in both human and *Drosophila* cells (Yang *et al*. 2014; Kemp *et al*. 2021). However, even these critical HLB proteins are poorly conserved at the sequence level, and there is little indication that Mxc interacts directly with DNA (Terzo *et al*. 2015; Kaya-Okur *et al*. 2019; Kemp *et al*. 2021). Therefore, Mxc and FLASH are unlikely to be the first factors that identify the *Drosophila* zygotic histone genes for unique regulation during development.

Mechanisms of histone gene regulation are well studied in *D. melanogaster*, as the histone genes reside at a single locus (Günesdogan *et al*. 2014; Bongartz and Schloissnig 2019) and histone array transgenes attract HLB factors (Salzler *et al*. 2013). Similar manipulative studies in mammalian systems are comparatively much more difficult (Sankar *et al*. 2022) due to the dispersion of histone genes across two chromosomes and megabases of sequence (Marzluff *et al*. 2002). In *D. melanogaster*, the histone locus is identified by the zinc-finger protein CLAMP, which interacts with long, perfect GA-repeat *cis* elements in the *H3/H4* promoter (Salzler *et al*. 2013; Rieder *et al*. 2017). The CLAMP-histone locus interaction promotes recruitment of Mxc and other HLB proteins (Rieder *et al*. 2017)(L. Hodkinson, observation). Removing the GA-repeat *cis* elements from histone array transgenes abrogates the ability of the transgene to recruit HLB-specific factors (Rieder *et al*. 2017)(L. Hodkinson, observation). At the endogenous locus, CLAMP increases histone locus chromatin accessibility and promotes histone gene expression of all five replication-dependent genes (Rieder *et al*. 2017). While there are likely multiple redundant mechanisms of HLB formation (Koreski *et al*. 2020), CLAMP is the first known factor that directly interacts with histone locus DNA sequence to promote recognition of the histone genes and recruitment of HLB-specific factors.

However, CLAMP is not specific to the histone locus; it is also critical for *Drosophila* dosage compensation (Soruco *et al*. 2013; Soruco and Larschan 2014), which increases X-linked gene expression to equalize gene dosage of males to females and the X-chromosome to the autosomes. CLAMP targets the male X-chromosome at GA-rich sequences (MSL recognition elements; MREs (Alekseyenko *et al*. 2008)), increases male X-chromosome accessibility, and recruits the Male Specific Lethal complex (MSLc) (Kuzu *et al*. 2016; Larschan *et al*. 2017). MSLc spreads across the chromosome and deposits the activating H4K16ac mark to increase male X-linked gene expression (Conrad *et al*. 2012). In addition to its role in histone gene regulation and dosage compensation, CLAMP targets autosomal sites in males and females to increase promoter accessibility and transcriptional elongation (Urban *et al*. 2017). CLAMP is also a component of the Late Boundary Complex, which forms in late-stage *Drosophila* embryos and impacts both dosage compensation (Kaye *et al*. 2017) and insulation within the bithorax Hox gene complex (Wolle *et al*. 2015; Kyrchanova *et al*. 2019). With so many critical functions, it is not surprising that CLAMP is comparatively well conserved amongst Drosophilidae, although it is unique to insects (Kuzu *et al*. 2016).

We previously hypothesized that the well-conserved CLAMP factor provides a bridge between histone locus *cis* elements and poorly conserved locus-specific factors such as Mxc/NPAT (Rieder *et al*. 2017). Similarly, MSLc proteins are poorly conserved (Kuzu *et al*. 2016) and CLAMP may provide a conserved link to newly evolving sex chromosomes (Alekseyenko *et al*. 2013).

Based on the above observations, we were surprised to observe that the male-specific component of MSLc, MSL2, targets one of the two autosomal histone loci in *Drosophila virilis*. MSL2 does not target the single histone locus in other *Drosophila* species and MSL2 is largely confined to the male X-chromosome in *D. melanogaster* (Lucchesi and Kuroda 2015). We report that the critical GA-repeat *cis* element in the *D. melanogaster H3/H4* promoter is almost unrecognizable in *D. virilis*, and more closely resembles GA-rich X-linked MREs. Both *D. melanogaster* and *D. virilis* CLAMP recognize the *D. virilis H3/H4* promoter region *in vivo*. However, *in vitro*, CLAMP does not require the GA-rich element in the *D. virilis* promoter sequence.

Our observations suggest that the two *D. virilis* histone loci are differentially regulated, as previously documented in yeast (Norris and Osley 1987; Cross and Smith 1988) and sea urchin (Marzluff *et al*. 2006). Since MSL2 is only expressed in male *Drosophila*, MSL2 might contribute to sex-specific regulation of the histone genes. Our observations indicate that context-specific transcription factors such as CLAMP may not always completely differentiate between genomic locations, resulting in cross-talk between regions with extremely different regulatory requirements.

## Methods

### Drosophila strains

We used the following stocks, maintained on standard cornmeal/molasses food and raised at 18°C: *Drosophila melanogaster* (y[1]w[1118]; +;+;+), *Drosophila virilis* (National *Drosophila* Species Stock Center #15010-1051.88), *Drosophila pseudoobscura* (NDSSC #14011-0121.217), and *Drosophila willistoni* (NDSSC #14030-0811.15).

### Cloning and transgenesis

We engineered plasmids that include a 5kb histone array sequence consisting of the 5 replication-dependent histone genes and their relative promoters where the *histone4* and *histone2A* genes are FLAG-tagged (24 bp) at the N-termini (Salzler *et al*. 2013) (original plasmid gift of Drs. Robert Duronio and William Marzluff). We used geneblocks (IDT) carrying the desired changes to alter the sequence of the histone gene array using Gibson cloning. Transgenic sequences are detailed in **Supplemental Table 3**. We inserted all 1× histone array transgenes into the genome at the VK33 attP site on chromosome 3L (65B2) (Venken *et al*. 2006) using PhiC3-mediated integration (Genetivision).

### Electrophoretic Mobility Shift Assays (EMSAs)

We performed EMSAs after Aoki *et al*. (2008) with minor modifications.

#### Late embryo nuclear extracts

We prepared embryo extracts from 6-18 hour Oregon R embryos collected on apple juice plates and aged 6 hours at room temperature. We performed nuclear extract preparation as in (Aoki *et al*. 2008). We omitted the final dialysis step described in Aoki *et al*. and completed the extraction with the final concentration of KCl at 360 mM.

#### DNA probes

We made EMSA probes (sequences in **Supplemental Table 2**) using PCR using gblocks (IDT) as templates and the following primers: *D. melanogaster* sequences: H3H4p F1: CACAGCACGAAAGTCACTAAAGAAC, H3H4p R1: GTTTGAAAACACAATAAACGATCAGAGC; *D. virilis* sequences: virilis H3H4p F1: CACCACGAATGTCACTGAGG, virilis H3H4p R1: TGTTAAAAACACAATAATCGTGCGTC. We 5’ end labeled one pmol of probe with γ-32P-ATP (MP Biomedicals) using T4 polynucleotide kinase (New England BioLabs) in a 50 μl total reaction volume at 37°C for 1 hour. We used Sephadex G-50 fine gel (Amersham Biosciences) columns to separate free ATP from labeled probes. We adjusted the volume of the eluted sample to 100 μl using deionized water so that the final concentration of the probe was 10 fmol/μl.

#### Shifts

We performed 20 μl binding reactions consisting of 0.5 μl (5 fmol) of labeled probe in the following buffer: 25 mM Tris-Cl (pH 7.4), 100 mM KCl, 1 mM EDTA, 0.1 mM dithiothreitol, 0.1 mM PMSF, 0.3 mg/ml bovine serum albumin, 10% glycerol, 0.25 mg/ml poly(dI-dC)/poly(dI-dC). We added 1 μl of nuclear extract and incubated samples at room temperature for 30 minutes. We loaded samples onto a 4% acrylamide (mono/bis, 29:1)-0.5× TBE-2.5% glycerol slab gel. We performed electrophoresis at 4°C, 180 V for 3-4 hours using 0.5× TBE-2.5% glycerol as a running buffer. We dried gels and imaged using a Typhoon 9410 scanner and Image Gauge software.

#### Recombinant CLAMP EMSAs

We performed recombinant CLAMP EMSAs using full-length recombinant, purified CLAMP protein (61.8 kDa) (Duan *et al*. 2021). We used the LightShift Chemiluminescent EMSA Kit (Thermo Fisher #20148) and performed 20 μl binding reactions with 1 μl CLAMP (1 μM), 1 μl biotinylated probe (0.3 μg/μl) and 1 ug/ul poly(dI-dC)/poly(dI-dC). We incubated reactions for 25 min at room temperature, ran on a 6% nondenaturing polyacrylamide gel and electrophoresed at 100 V for 2 hr in 0.5 X TBE. We performed semi-wet gravity transfer using the TurboBlotter (Cytiva) for 4 hours at room temperature using 20X SSC transfer buffer. We visualized the blotusing the Nucleic Acid Detection Module Kit (Thermo Fisher #89880).

### Western blotting

We collected *D. melanogaster* and *D. virilis* embryos on standard grape juice plates for 16 hours and dechorinated on the plate in 100% bleach for 2 minutes. We washed embryos in 1X PBS and then lysed and ground in RIPA buffer + protease inhibitor (Roche #11697498001). We spun samples at 20,000g for 5 minutes and retained the supernatant; this was repeated twice. We diluted the resulting protein lysate in 6X Laemmli sample buffer and ran samples on a 4 - 20% Bolt Bis-Tris gel. We transferred samples to a PVDF membrane, which we blocked for 1 hour in 3% BSA in TBS-T. We incubated the membrane overnight at 4°C with primary antibody at the following concentrations: anti-MSL2^*mel*^ serum at 1:100 (gift from Dr. Mitzi Kuroda) and anti-β actin at 1:1000 (CST #8457S). We washed the membrane 3× 5 minutes in TBS-T and then incubated it with secondary antibody (LI-COR IRDye^®^ 800CW/680RD Goat anti-Rabbit IgG) at 1:10,000 in TBS-T + 0.01% SDS for 1 hour. We washed the membrane 3× 5 minutes in TBS-T and 1× 5 minutes TBS before imaging (Bio-Rad ChemiDoc).

### Immunofluorescence on polytene chromosomes

We performed polytene chromosomes immunostaining from salivary glands dissected from sexed third instar *Drosophila* larvae raised at 18°C on standard cornmeal/molasses food. We passed glands through fix 1 (4% formaldehyde, 1% Triton X-100, in 1× PBS) for 1 min, fix 2 (4% formaldehyde, 50% glacial acetic acid) for 2 min, and 1:2:3 solution (ratio of lactic acid:water:glacial acetic acid) for 5 min prior to squashing and spreading. We washed slides in 1X PBS, then in 1% Triton X-100 (diluted in 1X PBS), and blocked for one hour in .5% BSA diluted in 1X PBS. We then incubated slides with primary antibodies diluted in blocking solution (antibodies specifics below) overnight at 4° C in a dark, humid chamber. We washed slides in 1 X PBS and incubated with secondary antibody diluted in blocking solution (antibody specifics below) for two hours at room temperature. We mounted slides in Prolong Diamond anti-fade reagent with DAPI (ThermoFisher, P36961), and imaged chromosome spreads on a Zeiss Scope.A1 equipped with a Zeiss AxioCam using a 40×/0.75 plan neofluar objective using AxioVision software.

### Antibodies

We used primary antibodies at the following concentrations: guinea pig anti-Mxc (1:5000; gift from Drs. Robert Duronio and William Marzluff), rabbit anti-MSL2 (1:150; gift from Dr. Victoria Meller, originally from Dr. Ron Richmond), goat anti-MSL3 serum (1:500; gift from Dr. Erica Larschan, originally from Dr. Mitzi Kuroda). All primary antibodies are raised against the *D. melanogaster* forms of the proteins. We used AlexaFluor secondary antibodies (ThermoFisher Scientific) at a concentration of 1:1000: goat anti-guinea pig AF647 (A-21450), goat anti-rabbit AF488 (A-11008), donkey anti-goat AF488 (A-11055).

### Bioinformatics

We used the online platform Galaxy (usegalaxy.org) (Afgan *et al*. 2018) to map ChIP-seq datasets to the histone gene array. We used the following datasets from NCBI GEO: GSE165833 (Villa *et al*. 2021) and GSE133637 (Rieder *et al*. 2019). We mapped reads to a single copy of the histone gene array as in Mckay *et al*. (2015) and normalized when possible to available input samples. We visualized data using Integrative Genomics Viewer (IGV) (Robinson *et al*. 2011). We used a custom R script to combine replicates, when available. The script is deposited at https://github.com/rieder-lab/Omics-Replicate-Merger.

### Sequence annotation and alignments

We annotated the *D. virilis* genome assembly (Kim *et al*. 2022) using SnapGene by searching for histone protein conservation with *D. melanogaster. D. virilis* genome assembly: ASM798932v2. *D. melanogaster* histone array sequences from (McKay *et al*. 2015). *D. virilis* histone array sequences from UCSC genome browser (http://genome.ucsc.edu) Aug. 2005 (Agencourt prelim/droVir2) release, scaffold_13047, range 1568499-1650698. *D. virilis* histone array sequences from (DDBJ accession no. AB249651) (Nakashima *et al*. 2016). We aligned sequences using Coffee (Erb *et al*. 2012) and SnapGene.

## Results

### Sequence differences between D. virilis histone loci

*Drosophila virilis* diverged from *D. melanogaster* ~ 40 million years ago. Instead of one histone locus, as in *D. melanogaster*, it carries two: a major locus on Chromosome II (25F) and a minor locus on Chromosome IV (43C)(Schienman *et al*. 1998). Schienman *et al*. (1998) performed Southern blot analysis and concluded that the major locus includes 25-30 arrays of both a quintet organization *(H2a, H2b, H3, H4*, and *H1*) and a quartet lacking *H1*, while the minor locus includes 6-8 quintet arrays. Kim *et al*. (2022) recently assembled 101 drosophilid genomes using PacBio long-read sequencing. We annotated the *D. virilis* histone loci from the Kim *et al*. assembly and determined that the major locus includes 5 quintet arrays and 27 quartet arrays, as well as many interrupted arrays and gene fragments. The minor locus includes 5 regularly spaced quintet arrays.

The ~100 regularly spaced quintet histone gene arrays in *D. melanogaster* are nearly identical in sequence (Bongartz and Schloissnig 2019). For example, the core *H3* genes vary at only two silent site locations in *D. melanogaster* (G231A, T408C) out of 411 total nt. However, we noticed substantial sequence differences between the same histone genes in *D. virilis:* five *H3* silent locations between the major and minor loci in *D. virilis*. Eight *H3* genes, distributed between the loci, include single nucleotide changes.

Similarity of the linker histone *H1* genes is more variable, compared to core histone genes; the human genome includes H1 subtypes H1.1-H1.5. Similarity of a subtype between species is higher than similarity of subtypes within a species (Di Liegro *et al*. 2018). By comparing the ~100 *D. melanogaster H1* genes, we found only two silent mutations (T312C, C462T) and two coding changes (D82E, V217I). However, we discovered that *D. virilis H1* genes include significant variation: 27 single nucleotide changes and 2 insertion/deletions (of 3 nucleotides each) across 753 nt. Eleven of the single nucleotide changes cause amino acid substitutions and the majority of sequence changes are locus-specific. Our observations suggest that replication-dependent *H1* subtypes across both loci could be present in this species.

### Both D. virilis histone loci are targeted by Mxc and CLAMP

The *D. virilis* histone loci differ in sequence, in contrast to the nearly identical sequences of the histone arrays at the single *D. melanogaster* locus. We therefore hypothesized that different factors target the major and minor loci. Mxc is an important HLB scaffolding protein (Hur *et al*. 2020; Kemp *et al*. 2021) that targets the *D. melanogaster* histone genes early during development (White *et al*. 2011). CLAMP is a non-histone-locus specific protein that interacts with GA-repeats in the *H3/H4* promoter, which promotes Mxc recruitment and HLB formation (Rieder *et al*. 2017). We previously observed that CLAMP targets histone loci in both *D. melanogaster* and *D. virilis* by polytene chromosome immunostaining (Rieder *et al*. 2017). We confirmed our previous observation by staining *D. virilis* polytene chromosomes with antibodies against *D. melanogaster* Mxc and CLAMP orthologs and observed that both Mxc and CLAMP target both *D. virilis* loci (**Figure 1A-B**).

**Figure 1:**
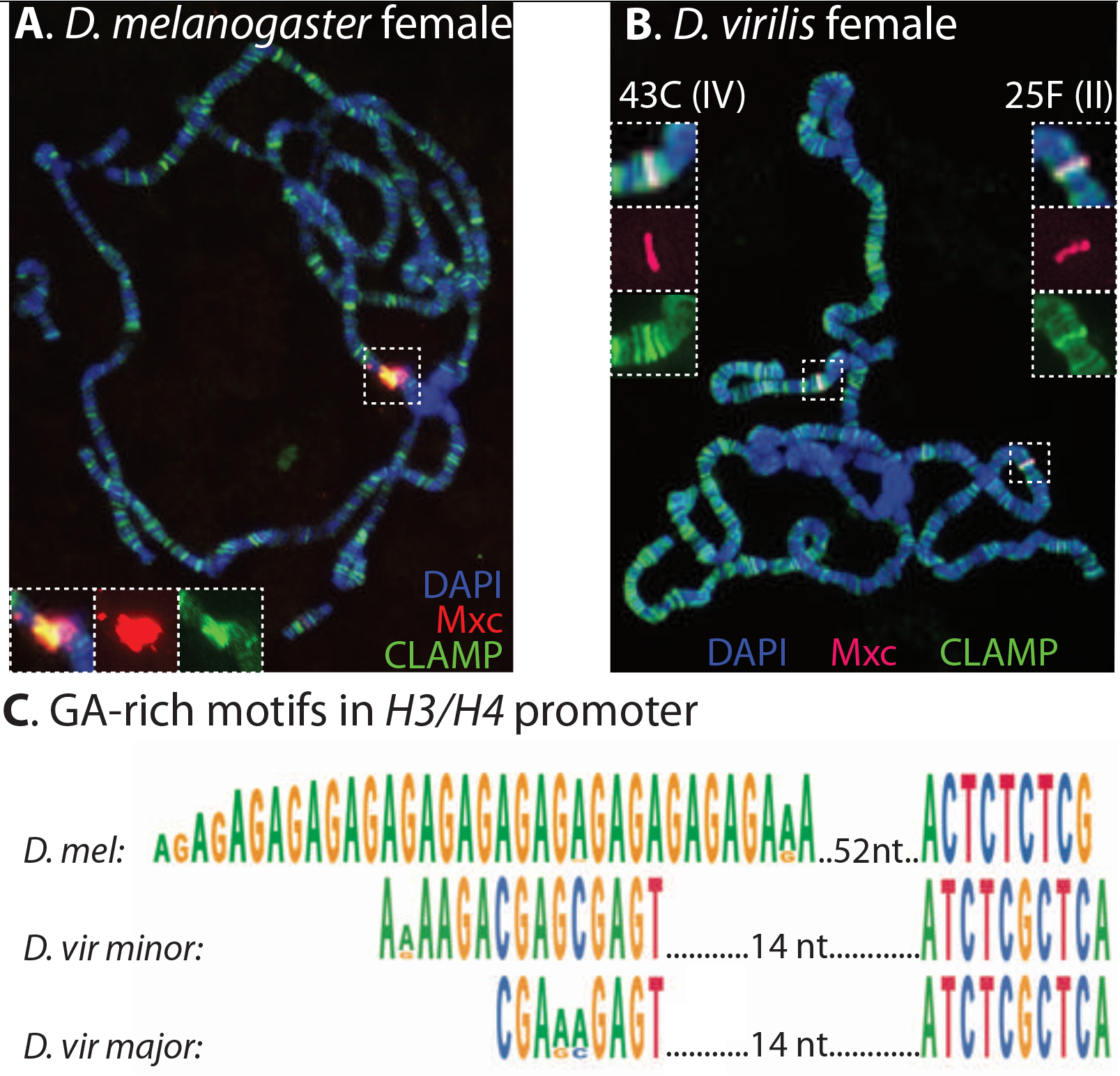
CLAMP and Mxc target the *D. virilis* histone loci. **A.** We confirmed previous results (Rieder *et al*. 2017) that CLAMP (green) targets the single histone locus in *D. melanogaster* by staining female larvae polytene chromosome spreads. Multi sex combs (Mxc; pink) is a core HLB protein that specifically targets the histone locus. **B.** CLAMP (green) also targets the two histone loci (colocalizing with Mxc) in *Drosophila virilis*. **C.** In *D. melanogaster (D. mel)*, CLAMP targets two long GA-repeats in the *H3/H4* promoter (Rieder *et al*. 2017). These GA-repeats are conserved but shorter and interrupted in *D. virilis* (*D. vir*).

CLAMP targets GA-rich motifs genome-wide in *D. melanogaster* early during development (Kuzu *et al*. 2016; Rieder *et al*. 2019) and facilitates several essential processes, including dosage compensation (Kuzu *et al*. 2016; Larschan *et al*. 2017; Rieder *et al*. 2019) and transcriptional elongation (Urban *et al*. 2017). The GA-repeats found in the *D. melanogaster H3/H4* promoter are long and unbroken while they are shorter and interrupted in *D. virilis* (**Figure 1C**). The *D. virilis H3/H4 cis* elements therefore more closely resemble the interrupted, GA-rich MREs found on *Drosophila melanogaster* X-chromosomes, which are critical for dosage compensation (Alekseyenko *et al*. 2008; Villa *et al*. 2016). We therefore considered that CLAMP might be attracting MSLc to the major *D. virilis* histone locus.

### MSL2 targets the major D. virilis histone locus

MSLc is confined to the male X-chromosome in *D. melanogaster* (Lucchesi and Kuroda 2015), and this pattern is apparent when staining third instar larval polytene chromosomes (**Figure 2A-B**). MSLc does not co-localize with Mxc, which marks the single histone locus on chromosome 2L. We stained male wild-type *D. virilis* polytene chromosomes for Mxc (to mark the histone loci), and MSL2, the male-specific structural component of MSLc (Bashaw and Baker 1995; Kelley *et al*. 1997). We were surprised to observe that MSL2 specifically targets the male *D. virilis* major histone locus on Chromosome II but not the minor histone locus on Chromosome IV (**Figure 2C**).

**Figure 2:**
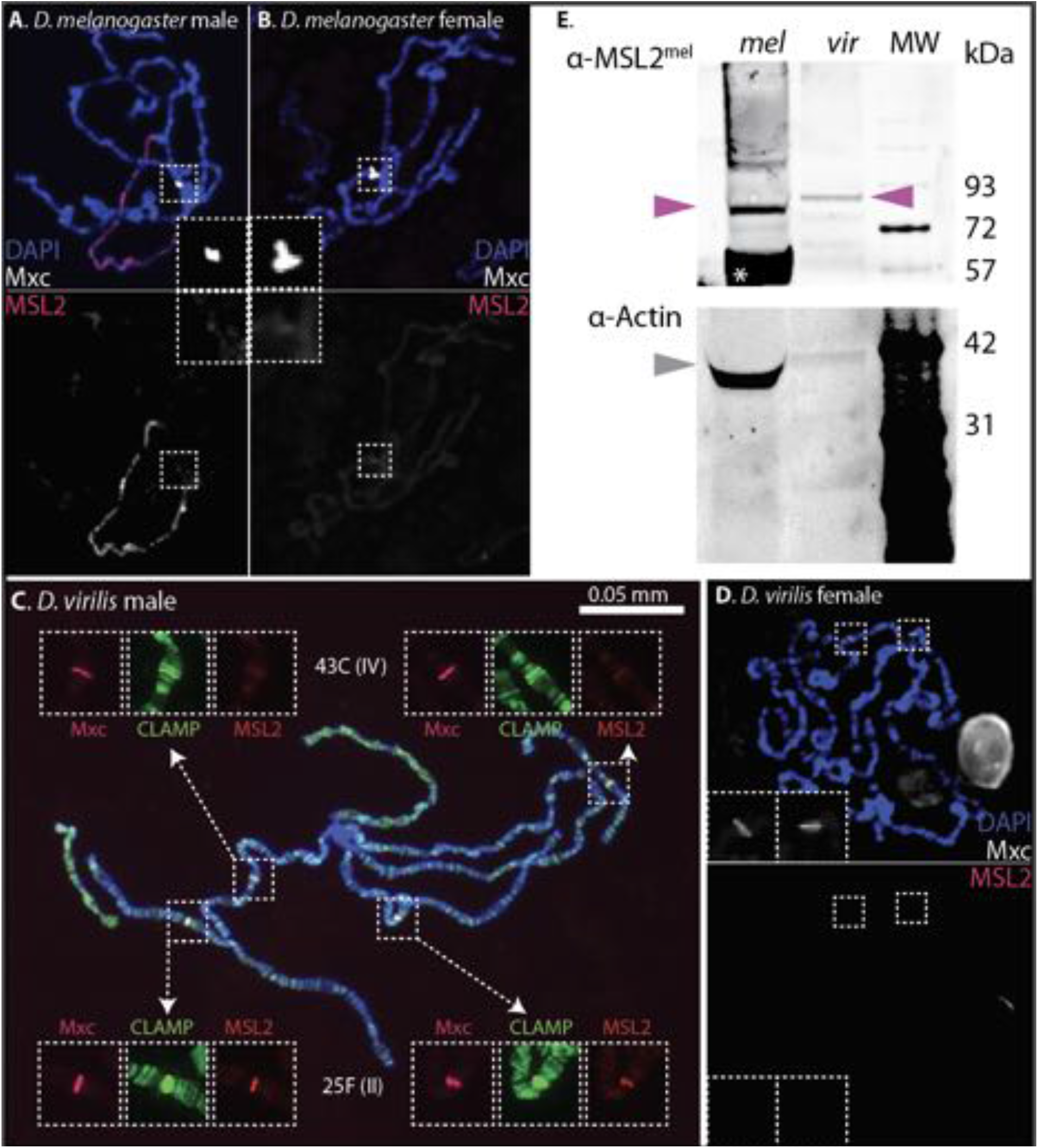
MSL2 targets only one histone locus in *D. virilis*. **A.** MSL2 (pink) targets the *D. melanogaster* male X-chromosome, but does not overlap with Mxc (white) at the histone locus. **B.** MSL2 is not present in female *D. melanogaster*. **C.** Two polytene chromosome spreads from *D. virilis* males show that MSL2 (red) signal overlaps with CLAMP (green) Mxc (pink) at the major *D. virilis* histone locus (25F) but not the minor locus (43C). **D.** MSL2 is not present in female *D. virilis*. **E.** The anti-*D. melanogaster* MSL2 antibody detects a ~81-85 kDa protein in both *D. melanogaster* and *D. virilis* embryo extracts, as well as a non-specific protein (*) present in both species. Samples are not concentration-normalized.

Our anti-MSL2 antibody was raised against *D. melanogaster* MSL2 sequence, and MSLc proteins are not well conserved even in Drosophilidae (Kuzu *et al*. 2016). Strangely, we did not observe male X-chromosome staining in *D. virilis* (**Figure 2C; Supplemental Table 1**), in contrast to prior work (Marín *et al*. 1996). Therefore, to confirm that our antibody is specific to MSL2 in both species, we performed western blotting. Our anti-MSL2^*mel*^ antibody recognized proteins around the correct predicted sizes: ~85 kDa in *D. melanogaster* and ~81 kDa in *D. virilis*, although the *virilis* ortholog may be modified and appears larger than the *melanogaster* ortholog (**Figure 2E**). Critically, we do not observe MSL2 staining at 25F in female *D. virilis* (**Figure 2D**), further indicating that the anti-MSL2^*mel*^ antibody is recognizing the *D. virilis* ortholog. These observations indicate that our antibody is specific to MSL2 and recognizes the protein in both species.

MSL2 is the structural component of MSLc and is usually found complexed with other members (Hallacli *et al*. 2012). However, MSL2 has some affinity for DNA sequence in both *D. melanogaster* and *D. virilis* (Villa *et al*. 2016, 2021) and CLAMP and MSL2 directly interact with each other through well-conserved domains (Tikhonova *et al*. 2022b), suggesting that CLAMP might recruit MSL2 outside of the complex. We therefore stained chromosome spreads for another MSLc member, MSL3. We did not observe recruitment of MSL3 to either of the histone loci in male *D. virilis* (**Figure 3**), indicating that MSL2 targets the major histone locus outside of its role in MSLc.

**Figure 3:**
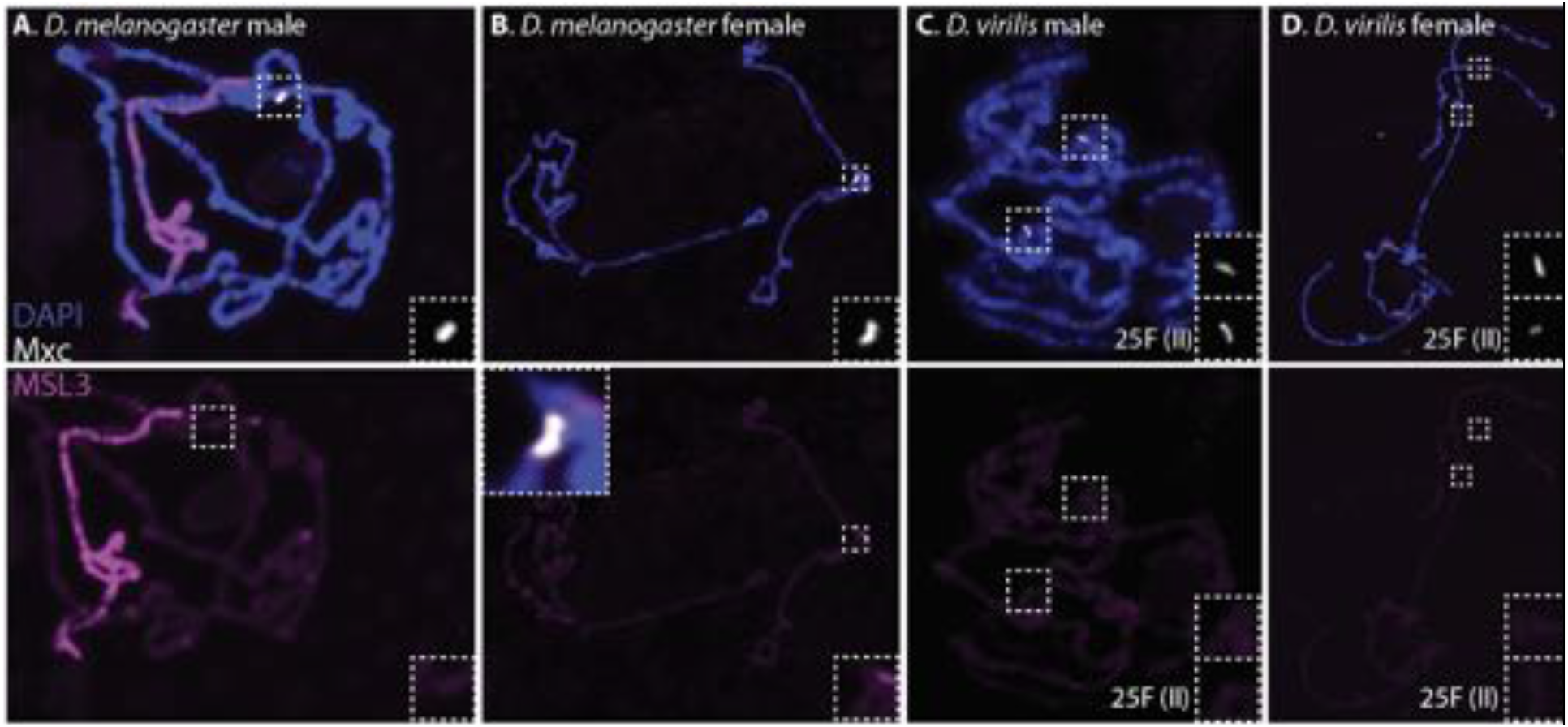
MSL3 does not target the histone locus in *Drosophila*. **A.** MSL3 (purple) targets the X-chromosome in *D. melanogaster* males, but does not colocalize with Mxc (white). **B.** MSL3 does not target loci on female *D. melanogaster* polytene chromosomes. **C.** MSL3 does not target the major histone locus (25F) in *D. virilis* males. **D.** MSL3 does not target loci on female *D. virilis* chromosomes.

We repeated our polytene experiments in *D. pseudoobscura* and *D. willistoni* (**Figure 4**), which both have a single histone locus. *D. melanogaster* and *D. pseudoobscura* diverged ~25 MYa, while *D. melanogaster* and *D. willistoni* diverged ~35 MYa (Powell 1997). We observed MSL2 specifically on the male X-chromosome in these species, indicating that MSLc targeting the histone genes is specific to *D. virilis* (**Supplemental Table 1**). These data suggest that MSLc specifically targets one of the two histone loci in *D. virilis*, a localization not observed in the other Drosophilids we investigated.

**Figure 4:**
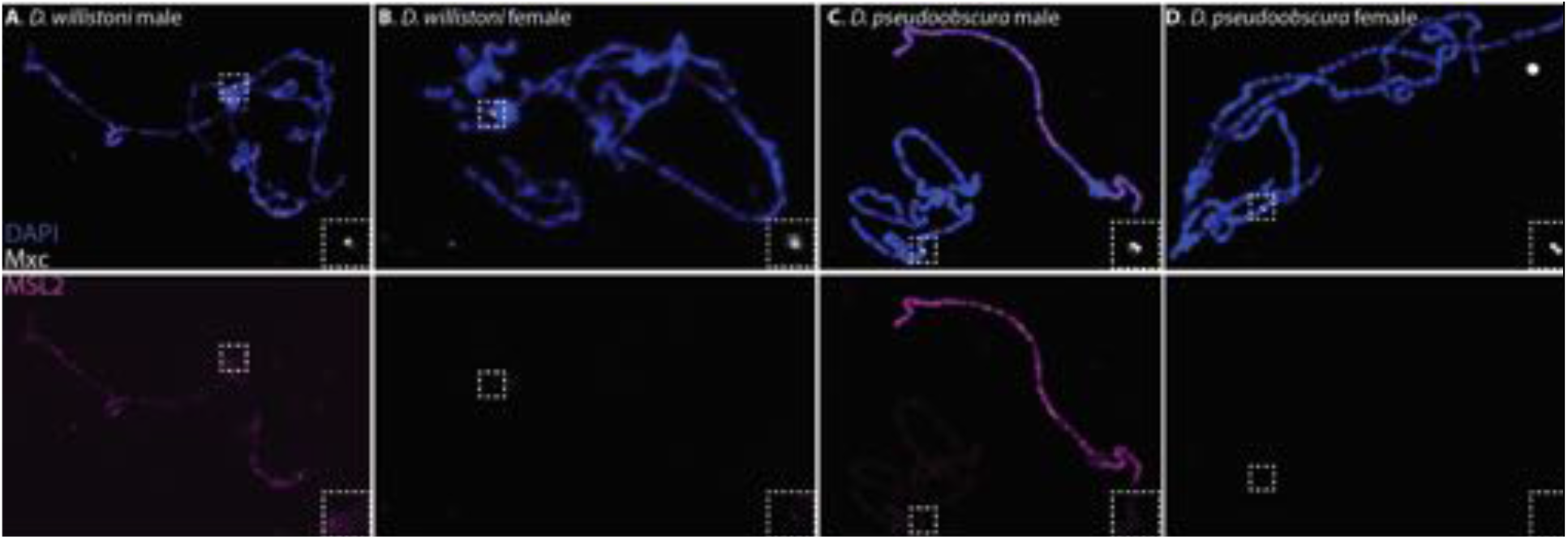
MSL2 does not target histone loci in other *Drosophila* species. MSL2 (purple) targets the X-chromosome in *D. willistoni* (**A**) and *D. pseudoobscura* males (**C**) but not females of either species (**B, D**). MSL2 does not co-localize with Mxc (white), which targets the single histone locus in both species.

### MSL2 does not directly interact with histone array sequence

CLAMP targets the zygotic *D. melanogaster* histone locus by nuclear cycle 10 (Rieder *et al*. 2017), just prior to detectable Mxc nuclear puncta (White *et al*. 2007) and zygotic histone gene expression in nuclear cycle 11 (Edgar and Schubiger 1986). Similarly, CLAMP targets loci genome-wide, including sites on the male X-chromosome, by nuclear cycle 11 (Rieder *et al*. 2019) and MSLc localizes to the male X-chromosome by nuclear cycle 14 (Rastelli *et al*. 1995). Although MSLc X-chromosome targeting requires CLAMP (Soruco *et al*. 2013), MSL2 has some ability to interact directly with DNA sequence (Villa *et al*. 2016, 2021; Tikhonova *et al*. 2019). Specifically, MSL2^*mel*^ identifies a subset of X-linked sequences called PionX sites (Villa *et al*. 2016), which often include GA-rich MRE elements. Villa et al. (2021) discovered that the *D. virilis* MSL2 ortholog is able to interact with DNA, but does not show the same specificity for X-linked sequences as the *D. melanogaster* ortholog. We hypothesized that MSL2 might be found specifically at the *H3/H4* promoter within the histone array, since that is where the CLAMP protein targets GA-repeats (**Figure 1C**) (Rieder *et al*. 2017), and CLAMP targets X-linked GA-rich MREs prior to MSLc in *D. melanogaster* (Rieder *et al*. 2019).

No studies have examined MSLc localization in *D. virilis* using genomics techniques. However, Villa *et al*. (2021) overexpressed GFP-tagged *D. melanogaster* (MSL2^*mel*^-GFP) and *D. virilis* (MSL2^*vir*^-GFP) MSL2 in cultured female *D. melanogaster* Kc cells and performed chromatin immunoprecipitation followed by sequencing (ChIP-seq). We mapped these datasets to a single *D. melanogaster* histone gene array (McKay *et al*. 2015). Since the array units in *D. melanogaster* are virtually identical in sequence (Bongartz and Schloissnig 2019), sequencing data is collapsed from ~100 arrays onto a single array. Neither *D. melanogaster* nor *D. virilis* MSL2 targets a sequence in the *D. melanogaster* histone gene array (**Supplemental Figure 1**). Importantly, we noticed that the control anti-GFP ChIP-seq dataset from untreated cells gives a sharp peak over the *H2a/H2b* promoter (**Supplemental Figure 2**), which is also found in other datasets. This peak in the control indicates that the GFP antibody interacts with sequences at the histone locus and likely elsewhere, confounding conclusions. In addition, we mapped MSL3 ChIP-seq datasets after MSL2 transfection (Villa *et al*. 2021) and did not observe enrichment over the *D. melanogaster* histone gene array (**Supplemental Figure 3**).

Finally, MSLc deposits the activating H4K16ac histone mark on the male X-chromosome (Gelbart *et al*. 2009). If MSLc is depositing H4K16ac at the major *D. virilis* histone locus, this post-translational modification could affect histone expression from the locus, specifically in males. In *D. melanogaster*, CLAMP targets sites across the X-chromosome, followed by MSLc and the appearance of H4K16ac by nuclear cycle 14 (Rieder *et al*. 2019). We mapped available H4K16ac ChIP-seq datasets from staged *D. melanogaster* male embryos (Rieder *et al*. 2019) to the histone gene array and did not observe H4K16ac enrichment (**Supplemental Figure 4**). However, this finding is not surprising given that we also do not observe MSLc at the *D. melanogaster* histone gene array by polytene chromosome immunostaining (**Figure 2A**) or by ChIP-seq (**Supplemental Figure 1**). We also mapped H4K16ac ChIP-seq datasets after MSL2^*mel*^ and MSL2^*vir*^ transfection (Villa *et al*. 2021) and did not observe H4K16ac enrichment over the *D. melanogaster* histone gene array (**Supplemental Figure 5**), which is not surprising since MSL2, but not other MSLc members, is present at the *D. virilis* major locus (**Figures 2, 3**).

We conclude that MSL2 does not directly interact with histone array sequence in either *D. melanogaster* or *D. virilis* and that other MSLc components are unlikely to be present at the major *D. virilis* histone locus.

### CLAMP does not require the GA-rich elements to interact with the *virilis H3/H4* promoter in vitro

It is difficult to assay CLAMP^*mel*^ recruitment to single histone array transgenes (L. Hodkinson, observation) since CLAMP is present at loci genome-wide (**Figure 1A-B**) (Urban *et al*.). It is also possible that CLAMP is present at the 1×His^*vir*^ transgene without interacting directly with DNA sequence, as was previously observed in histone array transgenes lacking GA-repeats (Koreski *et al*. 2020). We therefore turned to an *in vitro* approach to probe the interaction between CLAMP^*mel*^ and DNA sequence.

CLAMP is a member of the Late Boundary Complex (LBC) (Kaye *et al*. 2017) that forms in late embryogenesis (Wolle *et al*. 2015). We performed electrophoretic mobility shift assays using *D. melanogaster* late embryo extract and DNA probe sequences (**Supplemental Table 2**). We found that both *D. melanogaster* embryo extract, as well as recombinant full-length CLAMP^*mel*^ protein (Kuzu *et al*. 2016), shift both the wild-type *D. melanogaster* and *D. virilis H3/H4* sequences (**Figure 5**). The GA-rich *cis* elements in the *D. virilis* sequence are poorly conserved (**Figure 1C**) and there are other, short GA-repeats at other locations in the promoter. We therefore performed EMSAs using recombinant CLAMP^*mel*^ and 60 bp probes that tile the *D. virilis* promoter to confirm that CLAMP is targeting the region that contains the GA-rich *cis* element (**Supplemental Figure 6**).

**Figure 5:**
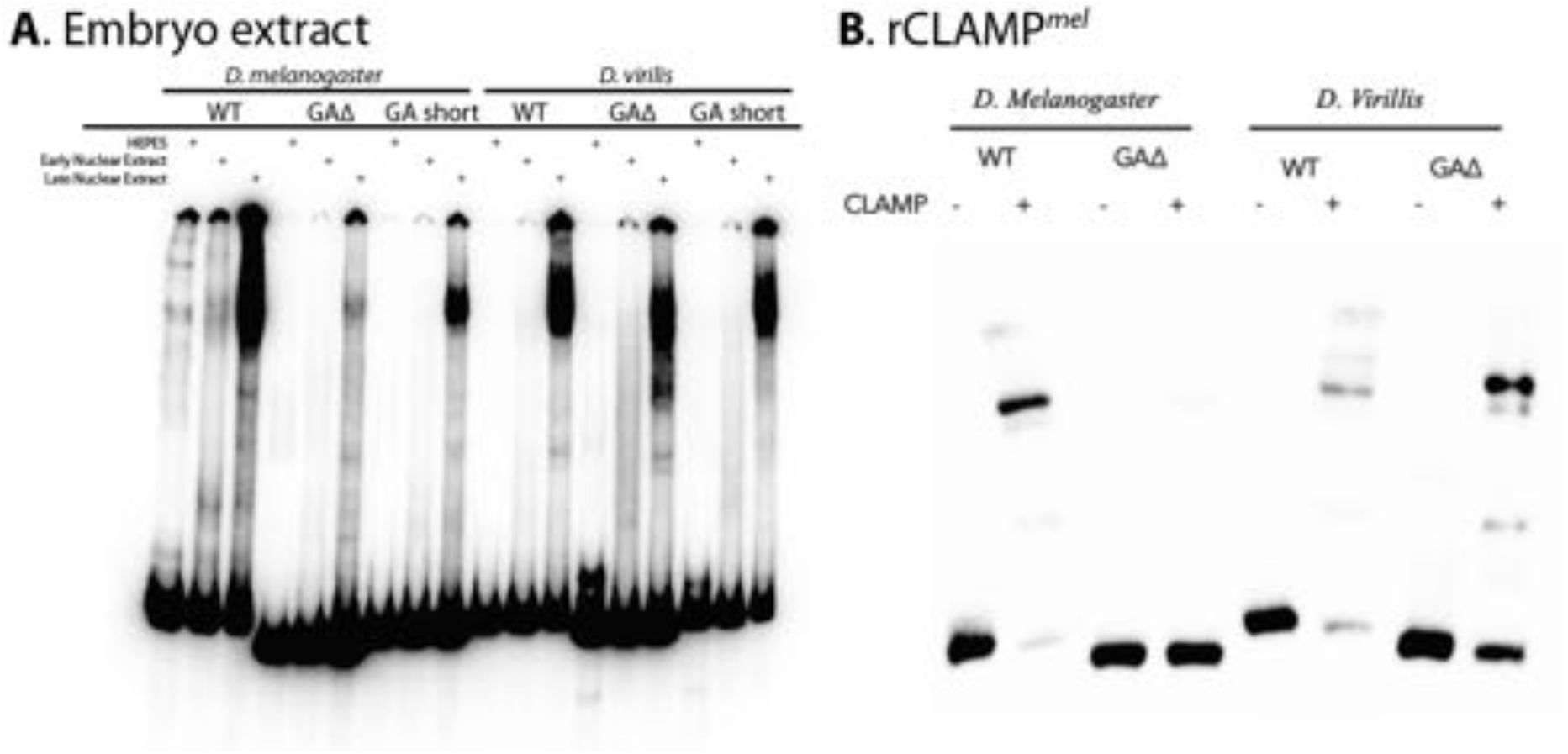
CLAMP binds the *D. virilis H3/H4* sequence *in vitro* and does not require the GA-rich elements. **A.** We shifted ^32^P-labeled dsDNA probes using protein extract from early (0-6hr) and late (6-18hr) *D. melanogaster* embryos. The wild-type (WT) *D. melanogaster H3/H4* sequence is shifted, but the shift is weaker when the GA-repeats are shortened (GA short). *D. melanogaster H3/H4* probe does not shift when the GA-repeats are removed (GAΔ). *D. virilis H3/H4* probes shift with *D. melanogaster* late embryo extract, even when the GA-rich element is removed. **B.** Recombinant full-length CLAMP^*mel*^ shifts the wild-type (WT) *D. melanogaster* and *D. virilis H3/H4* probes. Recombinant CLAMP continues to shift the *D. virilis* GAΔ probe. Probe sequences in **Supplemental Table 2.**

Deleting or shortening the GA-repeats in the *D. melanogaster* sequence compromises the shifting of the *melanogaster* probe. However, even deleting the GA-rich elements entirely from the *D. virilis* probe does not compromise shifting with *D. melanogaster* late embryo extract or recombinant full-length CLAMP (**Figure 5A-B**). Our *in vitro* observations suggest that CLAMP^*mel*^, and therefore likely CLAMP^*vir*^, is directly interacting with the *D. virilis H3/H4* promoter sequence. In addition, CLAMP^*mel*^, and therefore likely CLAMP^*vir*^, can target non-GA-repeat sequences in the *D. virilis H3/H4*.

### The D. virilis *H3/H4* promoter does not promote Mxc recruitment in D. melanogaster

Since CLAMP^*mel*^ is able to bind to the *D. virilis H3/H4* promoter sequence, we wondered if the *D. virilis* promoter might promote Mxc recruitment in *D. melanogaster*. It is difficult to manipulate the endogenous *D. melanogaster* histone locus, which includes ~100 nearly-identical histone gene arrays (McKay *et al*. 2015; Bongartz and Schloissnig 2019). However, wild-type *D. melanogaster* histone array transgenes recruit all tested HLB factors and express histone genes similar to the endogenous histone locus (Salzler *et al*. 2013; Rieder *et al*. 2017; Koreski *et al*. 2020) (1×His^WT^; **Figure 6A**). Histone array transgenes are therefore a powerful tool with which to interrogate DNA sequence contribution to histone locus identification.

**Figure 6:**
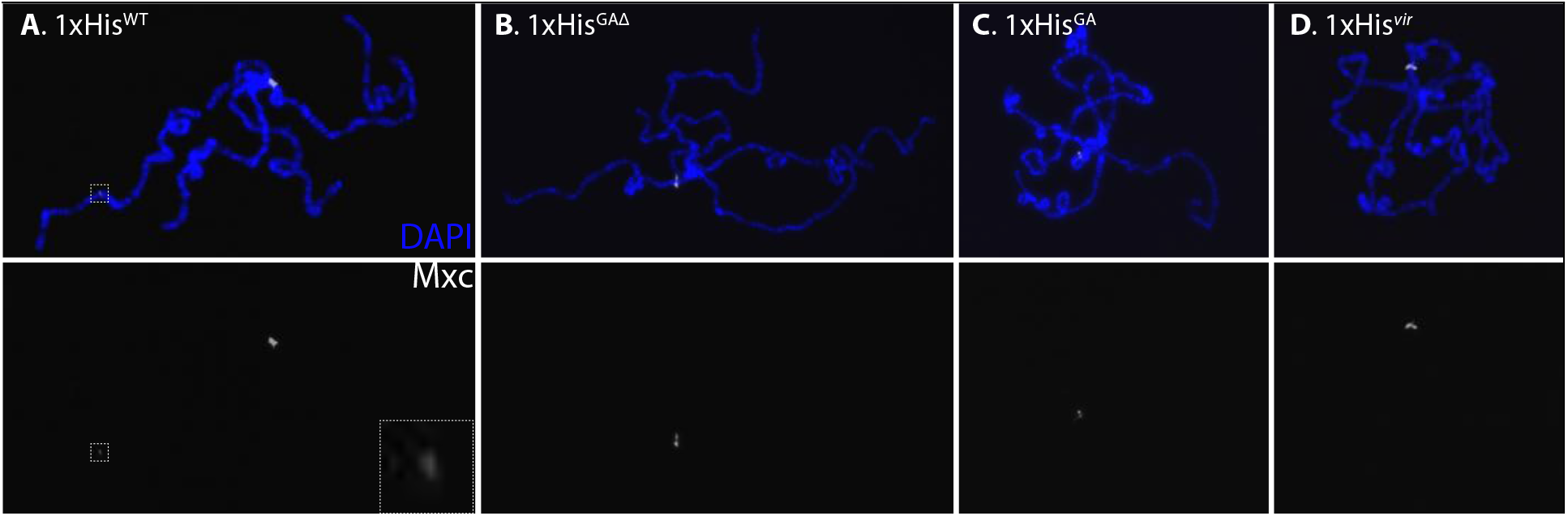
The *D. virilis H3/H4* promoter does not promote HLB formation in *D. melanogaster*. We performed polytene immunostaining for Mxc (white; bottom panels) in animals carrying 1× histone array transgenes. **A.** A wild-type transgene attracts Mxc (inset). **B.** Deleting the GA-repeats abrogates Mxc recruitment. **C.** Shortening the GA-repeats also abrogates Mxc recruitment to the transgene. **D.** Replacing the *H3/H4* promoter with that of *D. virilis* does not support Mxc recruitment. Transgene sequences in **Supplemental Table 3.**

As expected, deleting the GA-repeats from the *D. melanogaster H3/H4* promoter leads to failure to recruit the critical HLB scaffolding protein Mxc (1×His^GAΔ^; **Figure 6B**) (Rieder *et al*. 2017). We recently discovered that replacing the perfect GA-repeat sequence in a histone array transgene with X-linked CLAMP-binding *cis* elements abrogates HLB factor recruitment to the transgene (L. Hodkinson, observation), indicating the critical nature of the *cis* element sequence itself, rather than just the ability to recruit CLAMP.

The *D. virilis* GA-repeats in the *H3/H4* promoter are much shorter than that of *D. melanogaster* (**Figure 1C**) and they more closely resemble the X-linked GA-rich MREs involved in male dosage compensation (Alekseyenko *et al*. 2008; Villa *et al*. 2016). Although there are sequence differences between arrays, and even more differences between the major and minor loci, the GA-rich sequences are present in most arrays (**Supplemental Figure 7.**) We leveraged the transgenic histone array system to determine if the *D. virilis H3/H4* sequence is able to support Mxc recruitment in *D. melanogaster*.

We engineered a histone array transgene that replaces the majority of the *D. melanogaster H3/H4* promoter with a sequence from *D. virilis* (1×His^*vir*^ **Supplemental Table 3**). We also engineered a transgene with the *D. melanogaster* promoter sequence but shortened GA-repeats (1×His^GA^), using sequences similar to our *in vitro* gel shift assays (**Supplemental Table 2**). We discovered that neither transgene is able to recruit Mxc (**Figure 6C-D**). Since the *D. virilis* sequence is bound by CLAMP*^mel^ in vitro* (**Figure 5B**), our observations suggest that CLAMP targets the 1×His^*vir*^ transgene but is unable to recruit Mxc.

## Discussion

Here we show that MSL2 targets the major histone locus in *D. virilis*, but does not target the single histone locus in other *Drosophila* species. We propose that this is due to CLAMP interacting with the *D. virilis H3/H4* promoter sequence in a different manner than it does in *D. melanogaster*. The GA-rich element in the *D. virilis H3/H4* promoter more closely resembles GA-rich X-linked MREs (Alekseyenko *et al*. 2008) than it does the perfect, long GA-repeat of the *D. melanogaster H3/H4* promoter (Rieder *et al*. 2017). We recently discovered that X-linked MREs can drive MSL2 recruitment in the context of a transgenic histone gene array in *D. melanogaster* (L. Hodkinson, observation). However, our *in vitro* observations indicate that this element may even be dispensable for CLAMP interactions in *D. virilis*.

We previously observed that both *D. virilis* histone loci are targeted by CLAMP (Rieder *et al*. 2017). It is curious that we only observe MSL2 at the major locus. There are multiple suggested mechanisms to HLB formation, even in *D. melanogaster;* the GA-repeats are required in transgenic histone gene arrays for localization of CLAMP, Mxc, and other factors in *D. melanogaster*, as long as the transgene is in the background of the endogenous histone locus (Rieder *et al*. 2017). Transgenic arrays lacking the GA-repeats are targeted by Mxc only when the endogenous locus is deleted (Koreski *et al*. 2020). CLAMP is present in transgenic HLBs lacking GA-repeats by polytene chromosome immunostaining, but it does not interact with specific DNA sequences by ChIP-seq (Koreski *et al*. 2020). In addition to a zinc-finger domain, CLAMP contains a disordered prion-like domain (Kaye *et al*. 2018; Tikhonova *et al*. 2022a) that may facilitate dimerization and/or inclusion into the phase-separated HLB (Hur *et al*. 2020) likely through protein-protein interactions (Staller 2022).

This is not the first example of a degenerate *cis* elements facilitating a conserved interaction at the histone locus. In humans, octamer binding transcription factor 1 (Oct-1) controls S-phase H2B expression by targeting an 8 bp “octamer” element in the promoter (Zheng *et al*. 2003). Pdm-1/Nubbin is the Oct-1 counterpart in *Drosophila*, yet only cryptic octamer elements are found in both *H2B* and *H4* promoters and Pdm-1 influences expression of all core histone genes (Lee *et al*. 2010). Although humans and *Drosophila* share similar cell cycle needs for histone expression, histone octamer elements are conserved in vertebrates but not amongst *Drosophila* species. These observations led Lee *et al*. (2010) to suggest that invertebrates have greater tolerance for histone regulatory sequence flexibility, compared to vertebrates.

CLAMP is likely an evolutionary ancient protein that has been co-opted for multiple distinct functions (Kuzu *et al*. 2016; Rieder *et al*. 2017). CLAMP targets locations across the genome during very early embryogenesis. It targets the *D. melanogaster* histone locus around the same time as Mxc (Rieder *et al*. 2017; Kemp *et al*. 2021) and X-linked MREs prior to MSLc (Rieder *et al*. 2019). CLAMP and MSLc synergistically enrich each other’s occupancy on the male X-chromosome *in vivo* (Soruco *et al*. 2013; Soruco and Larschan 2014) and *in vitro* (Albig *et al*. 2019). We were surprised that we did not detect MSL2 on the male *D. virilis* X-chromosome, as was previously reported (Marín *et al*. 1996). Despite relatively low protein conservation (Copps *et al*. 1998; Kuzu *et al*. 2016), anti-MSL2^*mel*^ antibodies have long been used to assay other species. We discovered that MSL2^*vir*^ appears slightly larger than expected by western blot, which may indicate a post-translational modification on the majority of MSL2^*vir*^ that interferes with antibody-based detection.

However, MSLc components are not always confined to the X-chromosome. Males absent on the first (MOF) is an MSLc member that is present in other non-sex-specific complexes (Feller *et al*. 2012; Lam *et al*. 2012), while the Maleless helicase (MLE) member of the complex is expressed in both males and females and plays a role in RNA structure and splicing (Reenan *et al*. 2000). MSL2 is a core member of MSLc and is usually only present on the male X-chromosome. However, MSL2 mis-localizes to tandem repeats when the long non-coding *RNA on the X (roX)* components of the MSLc are missing in *D. melanogaster* (Figueiredo *et al*. 2014). Compromising the ability of MSL2 to interact with both CLAMP and DNA, via mutation of the MSL2 CLAMP-binding domain (CBD) and CXC domain, respectively, results in loss of complex X-chromosome specificity (Tikhonova *et al*. 2019). Both of these domains are well conserved between *D. melanogaster* and *D. virilis* (Villa *et al*. 2021). CLAMP and MSLc may search for genomic loci together as a “wolf pack” (Staller 2022) and have a higher affinity for sites resembling X-linked MREs. This model is supported by our recent observations that CLAMP targets loci genome-wide, and is followed by MSLc during very early *D. melanogaster* development, prior to MSLc X-chromosome specialization (Rieder *et al*. 2019).

While MSL2 is usually found complexed with other MSLc members (Hallacli *et al*. 2012), it retains some DNA binding ability in both *D. melanogaster (Villa et al. 2016)* and *D. virilis* (Villa *et al*. 2021). Further, White *et al*. (2011) identified MSL2, although not other MSLc proteins, in a proteomics screen for phosphorylated Mxc in cultured male *D. melanogaster* S2 cells. Mxc is phosphorylated by CyclinE/Cdk2, which activates *histone* gene expression and HLB phase separation (Wei *et al*. 2003; Hur *et al*. 2020). These observations provide evidence for the presence of MSL2 at histone loci, even sometimes in *D. melanogaster*.

Yet several lines of evidence argue against the presence of MSL2 at *D. melanogaster* histone genes. We do not observe MSL2 targeting the histone locus in *D. melanogaster in vivo* by polytene chromosome immunostaining or MSLc member ChIP-seq from multiple tissues. *D. melanogaster* polytene chromosome proximity ligation assays indicate that CLAMP only interacts with MSLc on the male X-chromosome (Lindehell *et al*. 2015). The above observations include both polytene chromosome immunostaining and analysis of sequencing datasets from cultured cells. Although the histone loci of polytene chromosomes appear to recruit all known HLB factors (Salzler *et al*. 2013; Rieder *et al*. 2017; Koreski *et al*. 2020), salivary gland nuclei undergo endoreplication without cell division and therefore might have unusual histone biogenesis requirements (Andreyeva *et al*. 2017). Cultured cells are asynchronous, and MSL2 might target the *D. melanogaster* histone locus at a specific cell cycle time point or in a subset of cell types, confounding results. Therefore, it is unlikely that the tissues examined here represent the histone regulatory needs of all tissues across developmental time.

It is curious that MSL2 targets only the major *D. virilis* histone locus on Chromosome II, which includes ~32 arrays of both the quintet and quartet organization (Schienman *et al*. 1998). The minor locus on Chromosome IV, which includes only ~5 quintet arrays, is targeted by CLAMP and Mxc but not MSL2. These observations indicate that the two histone loci may be differentially regulated in male *D. virilis*. There is no direct evidence that the ~100 nearly identical histone arrays in *D. melanogaster* are differentially regulated, however 100 copies are not required for viability; 12 transgenic arrays rescue viability when the endogenous locus is deleted (Günesdogan *et al*. 2014; McKay *et al*. 2015; Zhang *et al*. 2019).

It is not uncommon for histone genes to experience differential regulation. For example, the sea urchin genome carries three sets of histone genes: 2000 “early” α-histone genes are expressed in early embryogenesis, 35 “late” histone genes are expressed in somatic cells, and three histone genes are testes-specific (Marzluff *et al*. 2006). *Saccharomyces cerevisiae* mutants lacking one *H2a/H2b* unit *(TRT1)* cannot undergo mitosis, while those lacking the other unit *(TRT2)* have no dramatic phenotypes, indicating differential histone biogenesis from the two loci (Norris and Osley 1987; Cross and Smith 1988). The two histone loci in *D. virilis* may fulfill different developmental or cell cycle needs for histone production.

MSL2 is only expressed in XY *Drosophila; msl2* translation is repressed by Sex Lethal in XX individuals (Bashaw and Baker 1995; Kelley *et al*. 1997). The presence of MSL2 at the major histone locus in male, but not female, *D. virilis* indicates that histone genes might be differentially regulated between males and females. Although there is little current evidence of differential histone gene regulation between males and females, differences in sex chromosome size and chromatin content, and the existence of dosage compensation suggest that XX and XY individuals are likely to have different histone requirements. In addition, the requirement for maternal deposition of histone proteins (Horard and Loppin 2015) and the presence of maternal-effect *histone* gene-specific transcription factors such as *abnormal oocyte* (Berloco *et al*. 2001) indicate that males and females likely regulate histone loci differently, since female nurse cells must produce large amounts of histone mRNAs and proteins for egg deposition (Horard and Loppin 2015).

CLAMP participates in both male dosage compensation and histone biogenesis and therefore crosstalk between these gene regulatory networks (Friedlander *et al*. 2016) could occur in males but not in females. Factors are often shared between membraneless compartments (also called nuclear bodies). For example, Coilin is shared between Cajal and histone locus bodies at different developmental time points in *Drosophila* and *Xenopus* (Liu *et al*. 2006; Nizami *et al*. 2010). Nucleolin, fibrillarin, and other nucleolus factors are found in the Cajal body (Trinkle-Mulcahy and Sleeman 2017).

Overall, our results indicate that the two histone loci in *D. virilis* may be differentially regulated in males and females. The recruitment of MSL2 to the major *D. virilis* histone locus may be due to differential interactions between local DNA sequence and the CLAMP factor. Finally, CLAMP is shared between several compartments, which may lead to cross talk between gene regulatory networks.

## Acknowledgments

We thank John Ali for generating the original Galaxy pipeline. We thank Dr. Bob Duronio and Dr. William Marzluff for the anti-Mxc antibody, Dr. Victoria Meller (originally Dr. Ron Richmond) for the affinity purified anti-MSL2 antibody, Dr. Mitzi Kuroda for the anti-MSL2 serum, and Dr. Erica Larschan (originally Dr. Mitzi Kuroda) for the anti-MSL3 serum. We thank fellow group members Dr. Casey Schmidt, Tommy O’Haren, Greg Kimmerer, Eric Albanese, and Edgar Hsieh, who all supported this project.

## Funding

This work was supported by an Emory Summer Undergraduate Research Experience (SURE) Fellowship to MX, grants T32GM008490 and F31HD105452 to LJH, and grants K99HD092625, R00HD092625, and R35GM142724 (including S1) to LER.

## Conflict of interest

The authors declare no conflicts of interest.

## Notes

### Competing Interest Statement

The authors have declared no competing interest.

