## Supplemental Materials for "MSL2 targets histone genes in *Drosophila virilis*"

### Supplemental Figures and Tables:

**Supplemental Table 1: Summary of MSL protein localization from polytene chromosome immunostaining experiments. Related to Figures 2 and 4.**

| **Species** | **M/F** | **MSL2 at histone locus/loci?** | **MSL2 detected on X?** | **MSL3 at histone locus/loci?** | **MSL3 detected on X?** |
|---|---|---|---|---|---|
| *D. melanogaster* | M | No | **Yes** | No | **Yes** |
| | F | No | No | No | No |
| *D. virilis* | M | **Yes** | No | **Yes** | No |
| | F | No | No | No | No |
| *D. pseudoobscura* | M | No | **Yes** | Not tested | Not tested |
| | F | No | No | Not tested | Not tested |
| *D. willistoni* | M | No | **Yes** | Not tested | Not tested |
| | F | No | No | Not tested | Not tested |

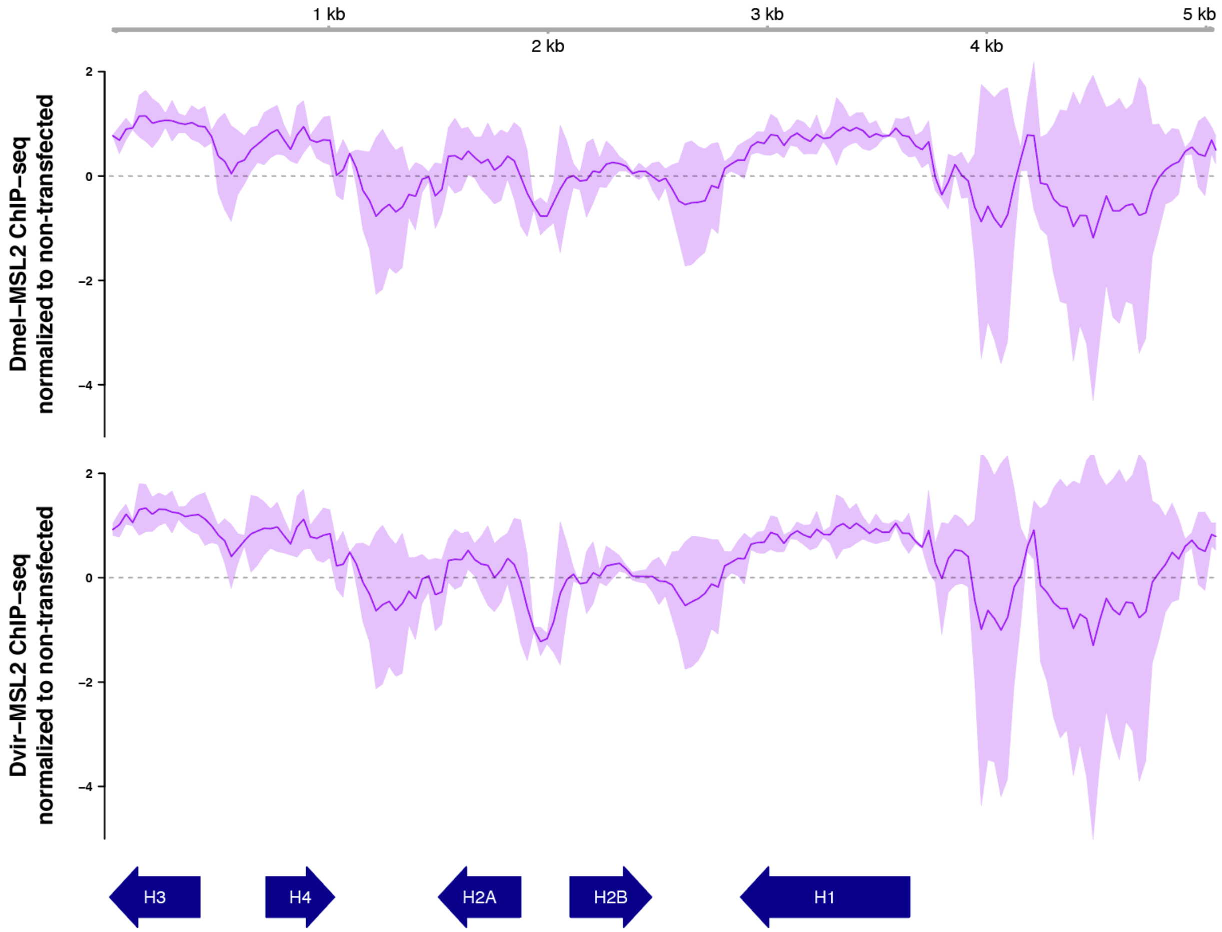


**Supplemental Figure 1: Ectopically expressed MSL2 from either species does not target the *D. melanogaster* histone gene array.** Villa *et al.* (2021) overexpressed GFP-tagged MSL2 from *D. melanogaster* and *D. virilis* in female *D. melanogaster* cell culture. They performed GFP ChIP-seq and we mapped their datasets to the *D. melanogaster* histone gene array, normalizing to the non-transfected control. Neither *D. melanogaster* MSL2 (top) nor *D. virilis* MSL2 (bottom) target a specific DNA sequence in the *D. melanogaster* histone gene array. The light purple indicates spread between two biological replicates, while the dark purple line indicates the average between replicates.

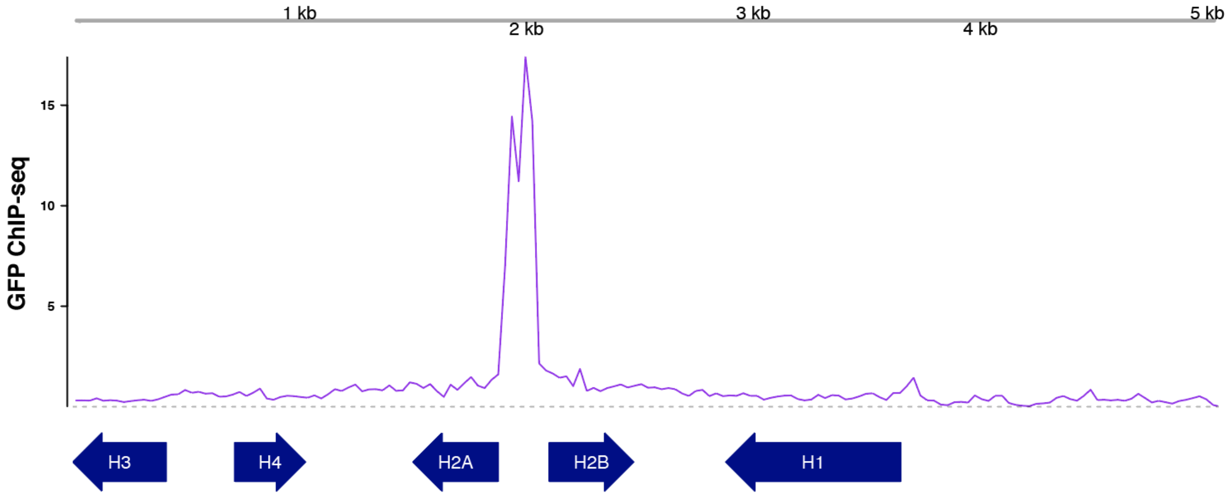


**Supplemental Figure 2: Anti-GFP ChIP-seq from non-transfected cells gives a peak in the *H2a/H2b* promoter.** Villa *et al.* (2021) overexpressed GFP-tagged MSL2 from *D. melanogaster* and *D. virilis* in female *D. melanogaster* cell culture. They performed GFP ChIP-seq in control, non-transfected cells and we mapped their datasets to the *D. melanogaster* histone gene array. We discovered a sharp peak in the *H2a/H2b* promoter, which is present in all anti-GFP ChIP-seq datasets.

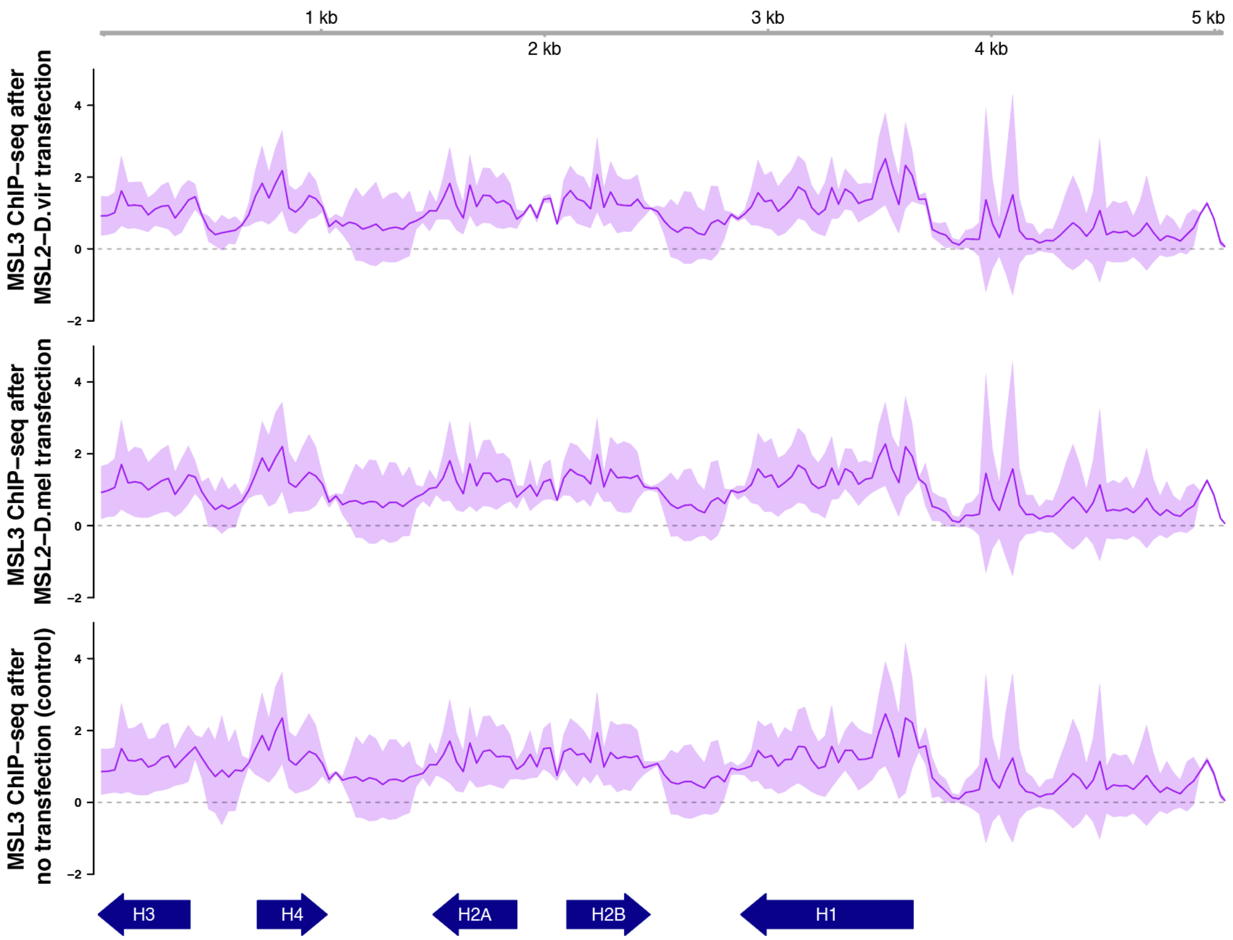


**Supplemental Figure 3: MSL3 does not target the *D. melanogaster* histone array after MSL2 transfection.** Villa *et al*. (2021) overexpressed MSL2 from *D. melanogaster* and *D. virilis* in female *D. melanogaster* cell culture. They performed MSL3 ChIP-seq and we mapped their datasets to the *D. melanogaster* histone gene array. MSL3 does not target the *D. melanogaster* histone gene array after transfection from either *D. virilis* MSL2 (top) nor *D. melanogaster* MSL2 (middle). These datasets look similar to MSL3 ChIP-seq from the non-transfected control (bottom). The light purple indicates spread between two biological replicates, while the dark purple line indicates the average between replicates.

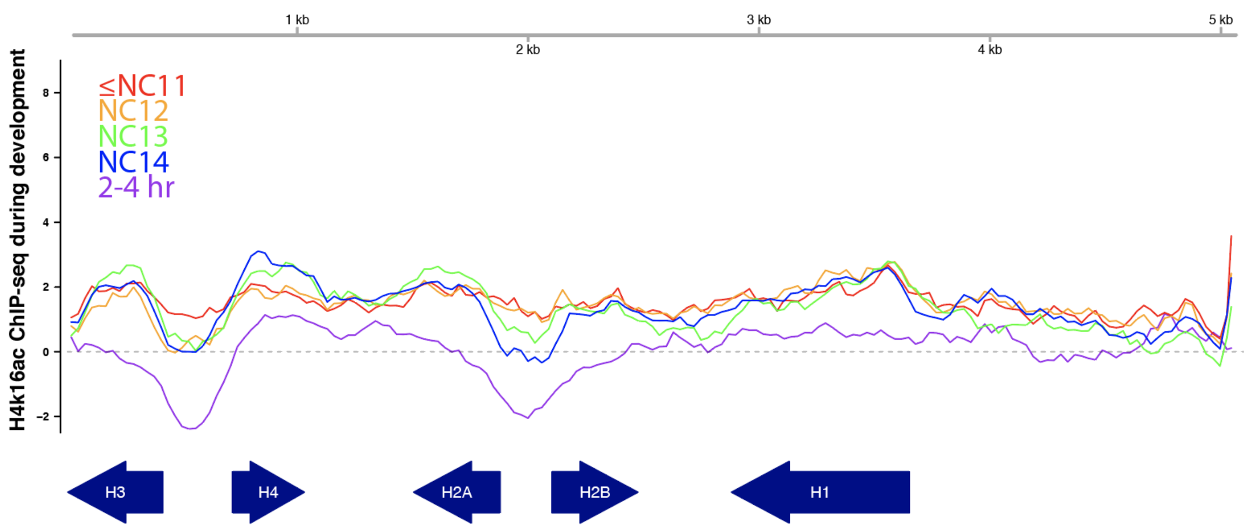


**Supplemental Figure 4: H4K16ac is not enriched over the *D. melanogaster* histone array during male *D. melanogaster* embryogenesis.** Rieder *et al*. (2019) performed male embryo H4K16ac ChIP-seq over a tight developmental time course (nuclear cycles = NC). We mapped single replicate datasets from this study to the *D. melanogaster* histone gene array, normalized to input samples. We observe no enrichment of H4K16ac over the *D. melanogaster* histone gene array.

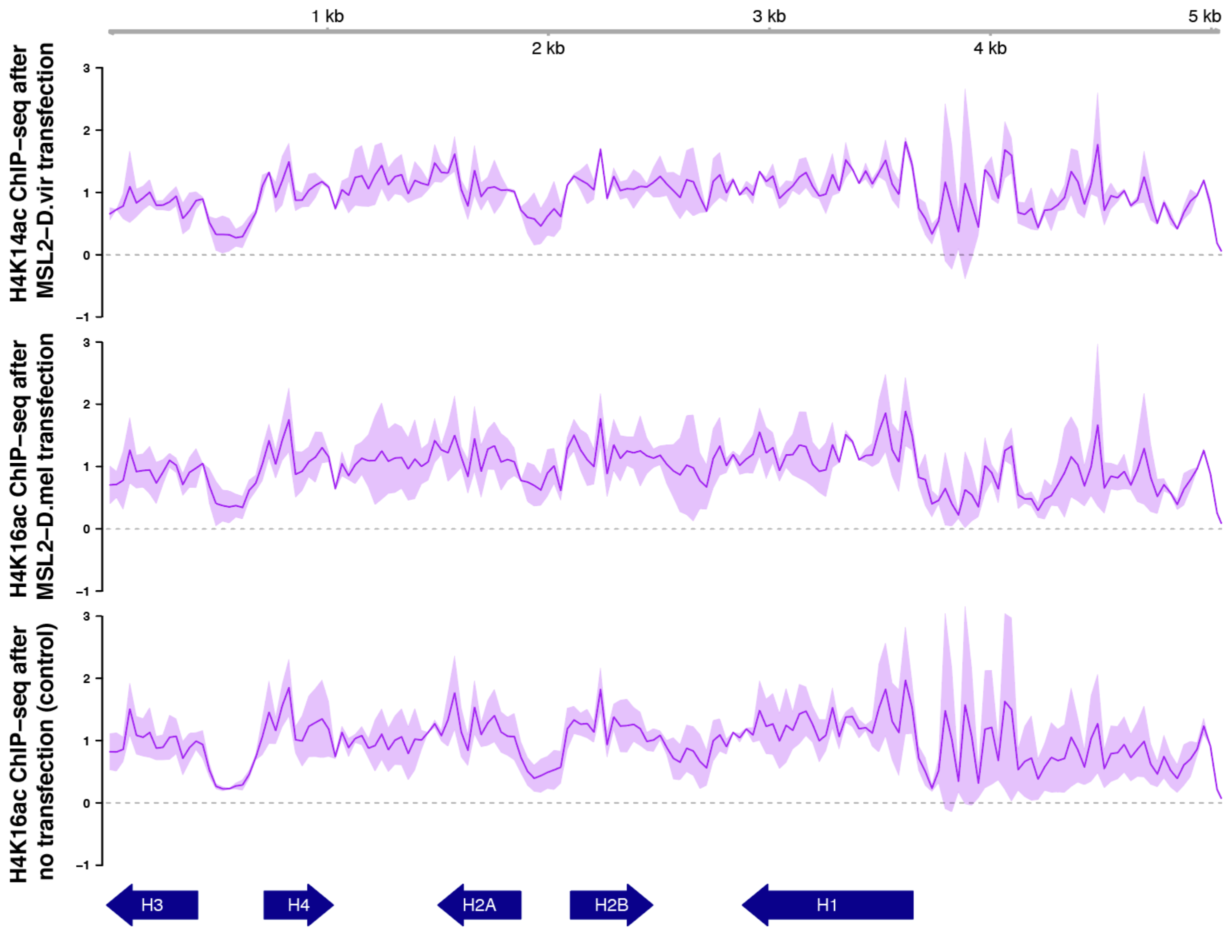


**Supplemental Figure 5: H4K16ac is not enriched over the *D. melanogaster* histone array after MSL2 transfection.** Villa *et al.* (2021) overexpressed MSL2 from *D. melanogaster* and *D. virilis* in female *D. melanogaster* cell culture. They performed H4K16ac ChIP-seq and we mapped their datasets to the *D. melanogaster* histone gene array. H4K16ac is not enriched over the *D. melanogaster* histone gene array after transfection from either *D. virilis* MSL2 (top) nor *D. melanogaster* MSL2 (middle). These datasets look similar to H4K16ac ChIP-seq from the non-transfected control (bottom). The light purple indicates spread between two biological replicates, while the dark purple line indicates the average between replicates.

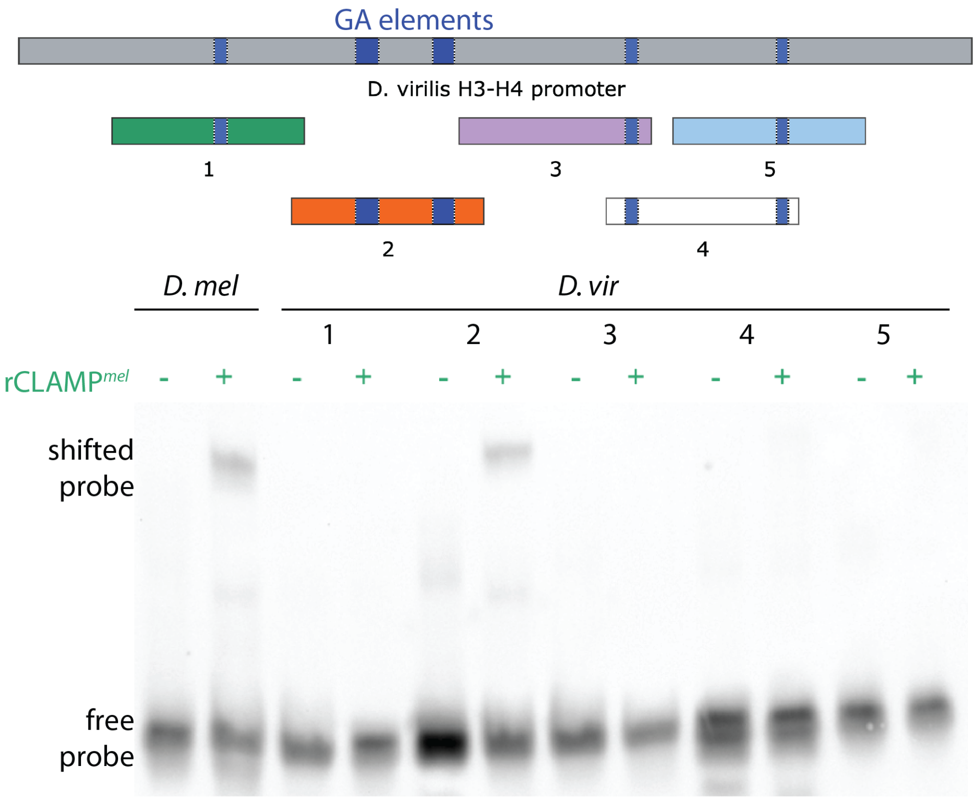


**Supplemental Figure 6: CLAMP interacts with the region containing the poorly conserved GA-rich *cis* elements in the *D. virilis* promoter.** The *D. virilis* promoter carries poorly conserved GA-rich elements (blue) as well as shorter GA-repeats (4 bp each; blue). We segmented the promoter into 60 bp probes (**Supplemental Table 2**) and performed EMSAs with recombinant CLAMP$^{mel}$. We confirmed that only probe 2 (orange), which carries the projected GA-rich *cis* elements, shifts with recombinant CLAMP.

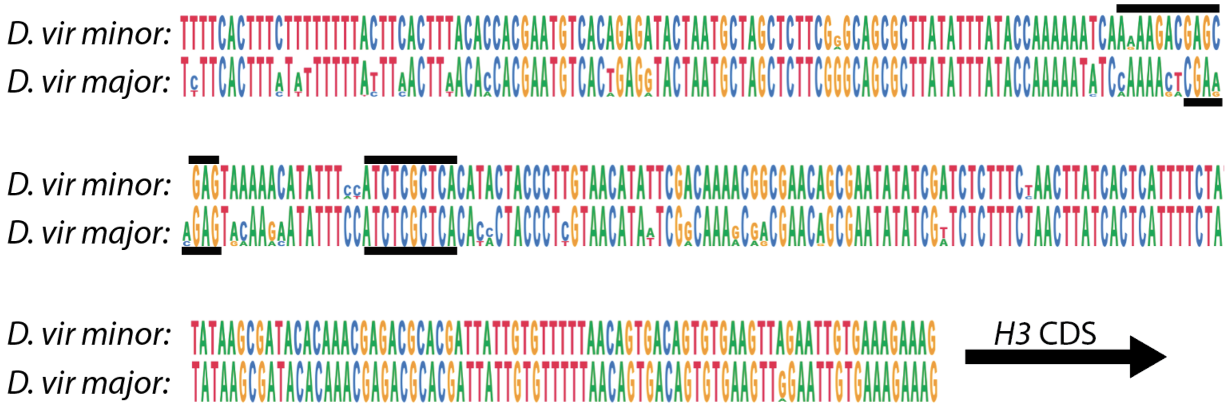


**Supplemental Figure 7: Conservation of the H3/H4 promoter within and between *D. virilis* loci.** We aligned sequences in SnapGene using Coffee (Erb *et al.* 2012). The minor (top) *D. virilis* locus includes five intact *H3/H4* promoters, while the major (bottom) includes 30 similar promoters. Conservation within loci is indicated by nucleotide height. GA-rich sequences shown in **Figure 1** are indicated by black bars. *H3* CDS represents the coding sequence, beginning with the start codon, of the *H3* gene.

**Supplemental Methods:**

**Supplemental Table 2: dsDNA EMSA probes. Related to Figures 5 and S7.**

| Probe (length) | Probe sequence |
|---|---|
| Wild-type *Dmel* **GA-repeats in bold** (226 bp; **Fig 6**) | CACAGCACGAAAGTCACTAAAGAACTAATTTCAACGTTTCTG TGTGCCCCTATTTATAGGTAAAACGACAAAAACCC**GAGAGA G**TACGAACGATATGTTCGTTCGCTTTTCGCTCGTCAAATGAA ATGGCCTCTGTTTT**TCTCTCTCTCTCTCTCTCTCT**TTCACCGT CCACGATTGCTATATAAGTAGGTAGCAAATGCTCTGATCGTT TATTGTGTTTTCAAAC |
| *Dmel* GA **GA-repeats in bold** (211 bp; **Fig 6**) | CACAGCACGAAAGTCACTAAAGAACTAATTTCAACGTTTCTG TGTGCCCCTATTTATAGGTAAAACGACAAAAACCCT**AGAGA**C TACGAACGATATGTTCGTTCGCTTTTCGCTCGTCAAATGAAA TGGCCTCTGTTTT**TCTCT**TTCACCGTCCACGATTGCTATATAA GTAGGTAGCAAATGCTCTGATCGTTTATTGTGTTTTCAAAC |
| *Dmel* GAΔ **Locations of deletions (X)** (198 bp; **Fig 6**) | CACAGCACGAAAGTCACTAAAGAACTAATTTCAACGTTTCTG TGTGCCCCTATTTATAGGTAAAACGACAAAAACCC**X**TACGAA CGATATGTTCGTTCGCTTTTCGCTCGTCAAATGAAATGGCCT CTGTTTT**X**TTCACCGTCCACGATTGCTATATAAGTAGGTAGC AAATGCTCTGATCGTTTATTGTGTTTTCAAAC |
| Wild-type *Dvir* **GA-repeats in bold** (235 bp; **Fig 6**) | CACCACGAATGTCACTGAGGTACTAATGCTAGCTCTTCGGG CAGCGCTTATATTTATACCAAAAACCAAAA**AGACGAG**CGAGT GAAAACATATTTCCATCTCGCTCACATACTACCCTTGTAACAT ATTCGACAAAACAGCGAACAGCGAATATATCGT**TCTCT**TTCT AACTTATCACTCATTTTCTATATAAGCGATACACAAACGAGAC GCACGATTATTGTGTTTTTAACA |
| *Dvir* GA **GA-repeats in bold** (231 bp; **Fig 6**) | CACCACGAATGTCACTGAGGTACTAATGCTAGCTCTTCGGG CAGCGCTTATATTTATACCAAAAACTCAAAAAGAC**AGAGA**TG AAAACATATTTCCAT**TCTCT**ACATACTACCCTTGTAACATATT CGACAAAACAGCGAACAGCGAATATATCGTTCTCTTTCTAAC TTATCACTCATTTTCTATATAAGCGATACACAAACGAGACGCA CGATTATTGTGTTTTTAACA |
| *Dvir* GAΔ **Locations of deletions (X)** (221 bp; **Fig 6**) | CACCACGAATGTCACTGAGGTACTAATGCTAGCTCTTCGGG CAGCGCTTATATTTATACCAAAAACTCAAAAAGAC**X**TGAAAAC ATATTTCCAT**X**ACATACTACCCTTGTAACATATTCGACAAAAC AGCGAACAGCGAATATATCGTTCTCTTTCTAACTTATCACTCA TTTTCTATATAAGCGATACACAAACGAGACGCACGATTATTGT GTTTTTAACA |
| Wild-type *Dmel* **GA-repeats in** | TGAAATGGCCTCTGTTTT**TCTCTCTCTCTCTCTCTCTCT**TT CACCGTCCACGATTGCT |

| | |
|---|---|
| **bold**<br>(60 bp; **Fig S7**) | |
| *Dvir* 1<br>**GA-repeats in bold**<br>(60 bp; **Fig S7**) | CACCACGAATGTCACTGAGGTACTAATGCTAG**CTCT**TCGGGCAGCGCTTATATTTATACC |
| *Dvir* 2<br>**GA-rich elements in bold**<br>(60 bp; **Fig S7**) | TACCAAAAACTCAAAAA**GACGAG**CGAGTGAAAACATATTTCCAT**CTCGCTC**ACATACTAC |
| *Dvir* 3<br>**GA-repeats in bold**<br>(60 bp; **Fig S7**) | CATACTACCCTTGTAACATATTCGACAAAACAGCGAACAGCGAATATATCGT**TCTCT**TTC |
| *Dvir* 4<br>**GA-repeats in bold**<br>(60 bp; **Fig S7**) | TATCGT**TCTCT**TTCTAACTTATCACTCATTTTCTATATAAGCGATACACAAAC**GAGA**CGC |
| *Dvir* 5<br>**GA-repeats in bold**<br>(60 bp; **Fig S7**) | TCACTCATTTTCTATATAAGCGATACACAAAC**GAGA**CGCACGATTATTGTGTTTTTAACA |

**Supplemental Table 3: *H3/H4* promoter sequences from *D. melanogaster* histone array transgenes. Related to Figure 6.**

| Transgene | Promoter sequence (between *H4* and *H3* start codons) |
|---|---|
| 1xHis$^{WT}$<br>(Hodkinson et al.)<br>**GA-repeats in bold**<br><br>(239 bp) | TTTTCACTGTTCTATACTATTATACACGCACAGCACGAAAGTC<br>ACTAAAGAACTAATTTCAACGTTTCTGTGTGCCCCTATTTATA<br>GGTAAAACGACAAAAACCC**GAGAGAG**TACGAACGATATGTT<br>CGTTCGCTTTTCGCTCGTCAAATGAAATGGCCTCTGTTTT**TCT**<br>**CTCTCTCTCTCTCTCTCT**TTCACCGTCCACGATTGCTATATAA<br>GTAGGTAGCAAATGCTCTGATCGTTT |
| 1xHis$^{GA\Delta}$<br>(Hodkinson et al.)<br>**Locations of deletions (X)**<br><br>(211 bp) | TTTTCACTGTTCTATACTATTATACACGCACAGCACGAAAGTC<br>ACTAAAGAACTAATTTCAACGTTTCTGTGTGCCCCTATTTATA<br>GGTAAAACGACAAAAACCC**X**TACGAACGATATGTTCGTTCGC<br>TTTTCGCTCGTCAAATGAAATGGCCTCTGTTTT**X**TTCACCGTC<br>CACGATTGCTATATAAGTAGGTAGCAAATGCTCTGATCGTTT |
| 1xHis$^{GA}$<br>**GA-repeats in bold**<br><br>(222 bp) | TTTTCACTGTTCTATACTATTATACACGCACAGCACGAAAGTC<br>ACTAAAGAACTAATTTCAACGTTTCTGTGTGCCCCTATTTATA<br>GGTAAAACGACAAAAACCCT**AGAGA**TACGAACGATATGTTC<br>GTTCGCTTTTCGCTCGTCAAATGAAATGGCCTCTGTTTT**TCTC**<br>**T**TTCACCGTCCACGATTGCTATATAAGTAGGTAGCAAATGCT<br>CTGATCGTTT |
| 1xHis$^{vir}$<br>**GA-rich elements in bold**<br><br>(297 bp) | TTTTCACTTTATATTTTTTTTTAACTTAACACCACGAATGTCAC<br>TGAGGTACTAATGCTAGCTCTTCGGGCAGCGCTTATATTTAT<br>ACCAAAAACTCAAAA**AGACGAGCGAG**TGAAAACATATTTCCA<br>**TCTCGCTC**ACATACTACCCTTGTAACATATTCGACAAAACAG<br>CGAACAGCGAATATATCGTTCTCTTTCTAACTTATCACTCATT<br>TTCTATATAAGCGATACACAAACGAGACGCACGATTATTGTG<br>TTTTTAACAGTGACAGTGTGAAGTTGGAATTGTGAAAGAAAG |
